# Mutational Robustness Predicts Protein Dynamics Across Natural and Designed Proteins

**DOI:** 10.64898/2026.03.19.713008

**Authors:** Or Zuk

## Abstract

Mutationally sensitive residues, where substitutions cause large stability changes, should also be dynamically rigid, because both properties arise from tight local packing and extensive contact networks. We test this prediction by defining a per-residue mutational robustness index, the standard deviation of predicted ΔΔ*G* values across all 19 single-amino-acid substitutions (which we compute using the structure-conditioned predictor ThermoMPNN), and correlating it with molecular-dynamics RMSF, crystallographic B-factors, and NMR-derived order parameters across ∼2,000 natural proteins, ∼400 de novo designs, and 759 NMR-characterized proteins. Robustness predicts dynamics well (median within-protein |*ρ*| ≈ 0.6), approaching the AlphaFold2 pLDDT confidence score while providing complementary information: robustness explains additional variance beyond pLDDT on every dataset, with the largest gains on designed proteins. The correlation is equally strong on de novo designs that lack any evolutionary history, pointing to a biophysical effect rather than a proxy for sequence conservation. A multiple regression model using a full 20-dimensional ΔΔ*G* profile per residue further outperforms all scalar summaries, showing that the identity of the substituted amino acid encodes dynamical information not captured by single-number predictors. Case studies on individual proteins, including the Zika virus capsid where pLDDT fails almost entirely, show that robustness maps physically meaningful dynamical patterns onto three-dimensional structures. Mutational robustness is thus a physically interpretable probe of the local fitness landscape that complements structural confidence scores for predicting protein flexibility.

## 1 Introduction

Protein function depends on a spectrum of conformational motions, from picosecond bond vibrations to millisecond domain rearrangements [Henzler-Wildman and Kern, 2007]. Understanding and predicting which residues are flexible and which are rigid remains a central goal in structural biology.

The structure of the protein fitness landscape, i.e. how stability and function change under mutation, determines both the course of molecular evolution and the biophysical properties of individual residues. *Mutational robustness*, the degree to which a protein tolerates amino acid substitutions without loss of stability, is a key feature of this landscape. Two complementary theoretical frameworks connect robustness to protein evolution and dynamics.

At the *protein level*, Bloom et al. [2006] showed that thermodynamically stable proteins tolerate a larger fraction of random mutations without losing their fold, making them more evolvable. Wagner [2008] further showed that robust systems can explore larger neutral networks in sequence space, resolving the apparent paradox between robustness and evolvability. This protein-level robustness depends on both native stability (Δ*G*) and the distribution of per-residue mutation effects (ΔΔ*G*).

At the *per-residue level*, Tokuriki and Tawfik [2009a] argued that conformational diversity, namely the ability of a protein to sample multiple states at specific sites, is itself linked to robustness, since flexible residues can accommodate mutations through alternative backbone conformations. This predicts a direct connection between local dynamics and local mutational tolerance that is independent of the protein’s overall stability.

These two frameworks converge for within-protein analysis. Since Δ*G* is a protein-wide constant, ranking residues within a single protein by mutational sensitivity depends entirely on their ΔΔ*G* values. Our central hypothesis is therefore: *mutationally sensitive residues should also be dynamically constrained (low RMSF), and mutationally robust residues should be more flexible (high RMSF).* A residue buried in a tight hydrophobic core has many stabilizing contacts, making it both intolerant to mutation and restricted in motion; a surface-exposed loop residue has fewer contacts, tolerates diverse substitutions, and fluctuates freely.

An alternative explanation is that both robustness and rigidity simply reflect evolutionary conservation: conserved sites are buried, mutationally sensitive, and rigid [Echave et al., 2015]. We disentangle the biophysical signal from the evolutionary one via two complementary tests: (i) testing the relationship on de novo designed proteins that lack any evolutionary record, and (ii) directly controlling for per-residue conservation scores from ConSurf [Ashkenazy et al., 2016] in the natural-protein analyses.

Recent computational and experimental advances now make systematic quantitative evaluation feasible. Large-scale MD simulation databases such as ATLAS [Vander Meersche et al., 2024] provide per-residue RMSF for thousands of proteins, while fast, accurate ΔΔ*G* predictors such as ThermoMPNN [Dieckhaus et al., 2024] and ESM-1v [Meier et al., 2021] enable genome-scale robustness computation. Furthermore, the PDB now contains hundreds of crystallized de novo designed proteins, enabling a direct comparison of the B-factor vs. robustness correlations between natural and designed proteins using the same experimental dynamics measure.

Supervised deep-learning tools such as PEGASUS [Vander Meersche et al., 2025b] can now predict MD-derived RMSF directly from protein language model embeddings, achieving per-residue Pearson correlations of ∼0.75 with MD ground truth; however, such models are trained directly on dynamics labels and do not reveal why a residue is flexible or rigid. De novo designed protein datasets with MD trajectories [Wolf et al., 2025] allow testing whether the relationship extends beyond natural proteins.

Here we test whether a per-residue mutational robustness index, defined as the standard deviation of predicted ΔΔ*G* across all 19 single-amino-acid substitutions, quantitatively predicts per-residue protein dynamics. We evaluate this index against MD-derived RMSF and crystallographic B-factors on both natural and designed proteins, and compare it to established baselines including the AlphaFold2 pLDDT confidence score [Jumper et al., 2021], which correlates moderately with MD flexibility but poorly with B-factors and NMR-derived dynamics gradations [Vander Meersche et al., 2025a, Gavalda-Garcia et al., 2025], and Solvent-Accessible Surface Area (SASA). Case studies on individual proteins where robustness substantially outperforms pLDDT illustrate the per-residue signal and its structural basis (Section 3.5).

## 2 Methods

### 2.1 Per-Residue Mutational Robustness Index

Let *S* = (*S*_1_*, S*_2_*, …, S_L_*) denote a protein sequence of length *L*, where each *S_i_* ∈ A = {A, C, D*, …,* Y} is one of the 20 standard amino acids. For each position *i*, we computed the predicted change in folding free energy ΔΔ*G_i_*_→*a*_ for every substitution *S_i_* → *a* where *a* ∈ A\{*S_i_*}. The **robustness index** at position *i* was defined as the standard deviation of the ΔΔ*G* distribution across all substitutions:

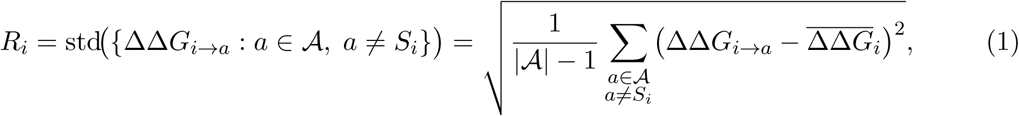

where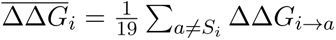 is the mean over the 19 substitutions at position *i*. A high *R_i_* indicates a mutationally sensitive position: one where some substitutions are highly destabilizing while others are tolerated, producing a wide spread of ΔΔ*G* values. A low *R_i_* indicates a robust position where all substitutions have similarly mild effects. Note that this per-residue, thermodynamic definition differs from the population-genetics concept of mutational robustness [Wagner, 2008, Bloom et al., 2006], which refers to the fraction of mutations a whole protein tolerates without loss of fold or function. Our index instead quantifies the *heterogeneity* of stability effects at a single site, and is used strictly for within-protein residue-level comparisons. This within-protein focus is justified because the native stability Δ*G* is a protein-wide constant: under a two-state folding model, a mutation is tolerated as long as ΔΔ*G_i_*_→*a*_ *<* |Δ*G*| (the protein still folds). When comparing residues within a single protein, Δ*G* cancels out and the ranking of mutational sensitivity is determined entirely by the ΔΔ*G* values.

We also evaluated alternative scalar summaries: the signed mean 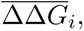 the absolute mean |ΔΔ*G*|, the maximum max |ΔΔ*G_i_*_→*a*_|, the fraction of destabilizing mutations (ΔΔ*G >* 1 kcal/mol), and the fraction of neutral mutations (|ΔΔ*G*| *<* 0.5 kcal/mol). The standard deviation isolates the heterogeneity of stability effects at each position, distinguishing sites where all mutations are moderately destabilizing from those with a mix of neutral and catastrophic substitutions. Empirical comparison is presented in Section 3.4.

We computed ΔΔ*G* using two predictors: **(i) ThermoMPNN** [Dieckhaus et al., 2024] is a structure-conditioned graph neural network trained on experimental ΔΔ*G* data from the MegaScale dataset [Tsuboyama et al., 2023]; it takes the protein backbone structure and sequence as input. **(ii) ESM-1v** [Meier et al., 2021] is a protein language model that predicts mutation effects from sequence alone via masked marginal log-likelihoods, limited to proteins ≤ 1024 residues.

#### Structure preprocessing and residue matching

ThermoMPNN uses the ProteinMPNN backbone parser [Dauparas et al., 2022] to extract residue coordinates from PDB files. For **AT-LAS**, the MD pipeline had already converted modified residues to standard counterparts (e.g. MSE → MET) and removed non-protein atoms, so the parser-derived and canonical sequences agreed exactly. **BBFlow** structures contained only standard amino acids by construction. For **PDB de novo designs**, raw deposited coordinates could contain modified residues (HETATM entries) or expression-tag residues; we used the parser-derived sequence for all downstream analyses, ensuring consistent alignment of robustness, B-factor, and sequence annotations across the structurally resolved residues.

### 2.2 Dynamics Measures

We used two per-residue measures of protein dynamics. Let **r***_i_*(*t*) denote the C*_α_* atom position of residue *i* at time *t*, and 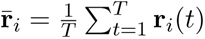 its time-averaged position.

#### Root mean square fluctuation (RMSF)

From an MD trajectory with *T* time frames:

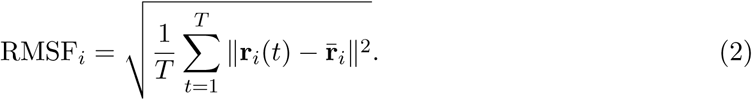

#### Crystallographic B-factor

The B-factor (Debye-Waller factor) is the experimental analogue from X-ray crystallography [Sun et al., 2019]:

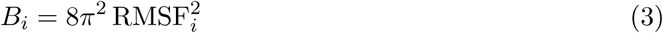

where the mean square displacement is now measured in the crystal lattice rather than in simulation. B-factors are noisier than MD-RMSF due to crystal packing effects and refinement artifacts, but provide an experimentally grounded dynamics measure.

### 2.3 Datasets

#### ATLAS (natural proteins)

We used the ATLAS database [Vander Meersche et al., 2024], which provides standardized all-atom MD simulations (3 × 100 ns replicates, CHARMM36m force field) for 1,938 monomeric protein chains from the PDB (444,240 residues), along with per-residue RMSF (averaged across replicates), AlphaFold2 pLDDT scores, and crystallographic B-factors. For the ESM-1v comparison only, 10 proteins with ≥ 1,024 residues were excluded due to the model’s context-length limit; all other analyses use the full set of 1,938 proteins.

#### BBFlow (de novo designed proteins, MD dynamics)

BBFlow [Wolf et al., 2025] is a generative model that produces protein backbone structures by learning from the conformational ensembles in the PDB; it is trained on backbone geometry (bond angles and dihedrals) rather than on sequence, and outputs diverse backbone folds that are subsequently threaded with ProteinMPNN [Dauparas et al., 2022] to obtain designable sequences. We analyzed 100 BBFlow-designed proteins, each with 3-replicate all-atom MD trajectories (same CHARMM36m protocol as ATLAS). Because no experimental structures exist, we computed per-residue pLDDT via ESMFold [Lin et al., 2023] and SASA from the design-model PDB using the Shrake-Rupley algorithm in mdtraj [McGibbon et al., 2015]. Total: 21,115 residues.

#### PDB de novo designs (designed proteins, B-factor dynamics)

To enable a direct natural-vs.-designed comparison using the same experimental dynamics measure, we assembled a dataset of de novo designed protein crystal structures from the PDB. We queried the RCSB PDB Search API with three full-text terms (“de novo designed protein”, “de novo protein design”, “computationally designed protein”), then applied quality filters matching the ATLAS criteria: X-ray diffraction only, resolution ≤ 2.0 ^Å^, chain length ≥ 38 residues, single protein entity, and at least one monomeric biological assembly; we excluded membrane proteins. Title-based filtering removed entries describing natural proteins with *designed ligands* (e.g. “inhibitor”, “in complex with”) unless the title also contained a positive design keyword (“de novo design”, “Rosetta”, “ProteinMPNN”, etc.); we similarly excluded structural-genomics entries.

After all filters, 318 proteins remained. We further excluded 12 entries that, upon inspection, turned out to be natural proteins, natural-scaffold variants, or had irreconcilable sequence mismatches rather than being genuine de novo designs (identified by cross-referencing against ConSurf-DB [Ashkenazy et al., 2016]: entries with ConSurf conservation scores whose PDB titles lacked any design-related keywords were flagged and manually verified). After this curation, 306 proteins remained (58,886 residues). We extracted per-residue C*_α_* B-factors from the deposited coordinates and computed per-residue pLDDT via ESMFold [Lin et al., 2023]. Of the 306 proteins, we successfully processed 290; 16 exceeded the GPU memory limit due to sequence length and were excluded from pLDDT-dependent analyses only (robustness and SASA analyses use all 306 proteins).

#### NMR-derived order parameters 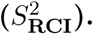

To validate the robustness-dynamics relationship against an experimental modality independent of MD simulation, we used the 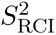 dataset of Gavalda-Garcia et al. [2025]. This dataset provides per-residue order parameters derived from NMR chemical shifts via the Random Coil Index method for 759 proteins (76,642 residues). 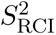 ranges from 0 (fully flexible) to 1 (fully rigid); we used 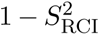 as the dynamics target so that high values indicate flexibility, matching the RMSF and B-factor convention. We obtained per-residue pLDDT from AlphaFold2 predicted structures and computed ThermoMPNN robustness from the same structures.

#### ConSurf evolutionary conservation scores

To disentangle the biophysical robustness signal from evolutionary conservation, we obtained per-residue conservation scores from ConSurf-DB [Ashkenazy et al., 2016] for the ATLAS proteins. ConSurf computes position-specific evolutionary rates using the Rate4Site algorithm, which estimates per-residue substitution rates from a multiple sequence alignment of homologs, corrected for the phylogenetic tree. Lower Rate4Site scores indicate higher conservation (slower evolutionary rate). Of the 1,938 ATLAS proteins, 1,847 (95%) had ConSurf scores available in ConSurf-DB, covering 427,362 residues. Conservation scores were used both as an independent predictor of dynamics (Table 1) and as a covariate in partial correlation analyses to test whether the robustness-dynamics signal survives controlling for evolutionary constraint (Supplementary Table S1). Conservation data are available only for the natural-protein datasets (ATLAS); designed proteins lack evolutionary homologs by construction.

**Table 1:**
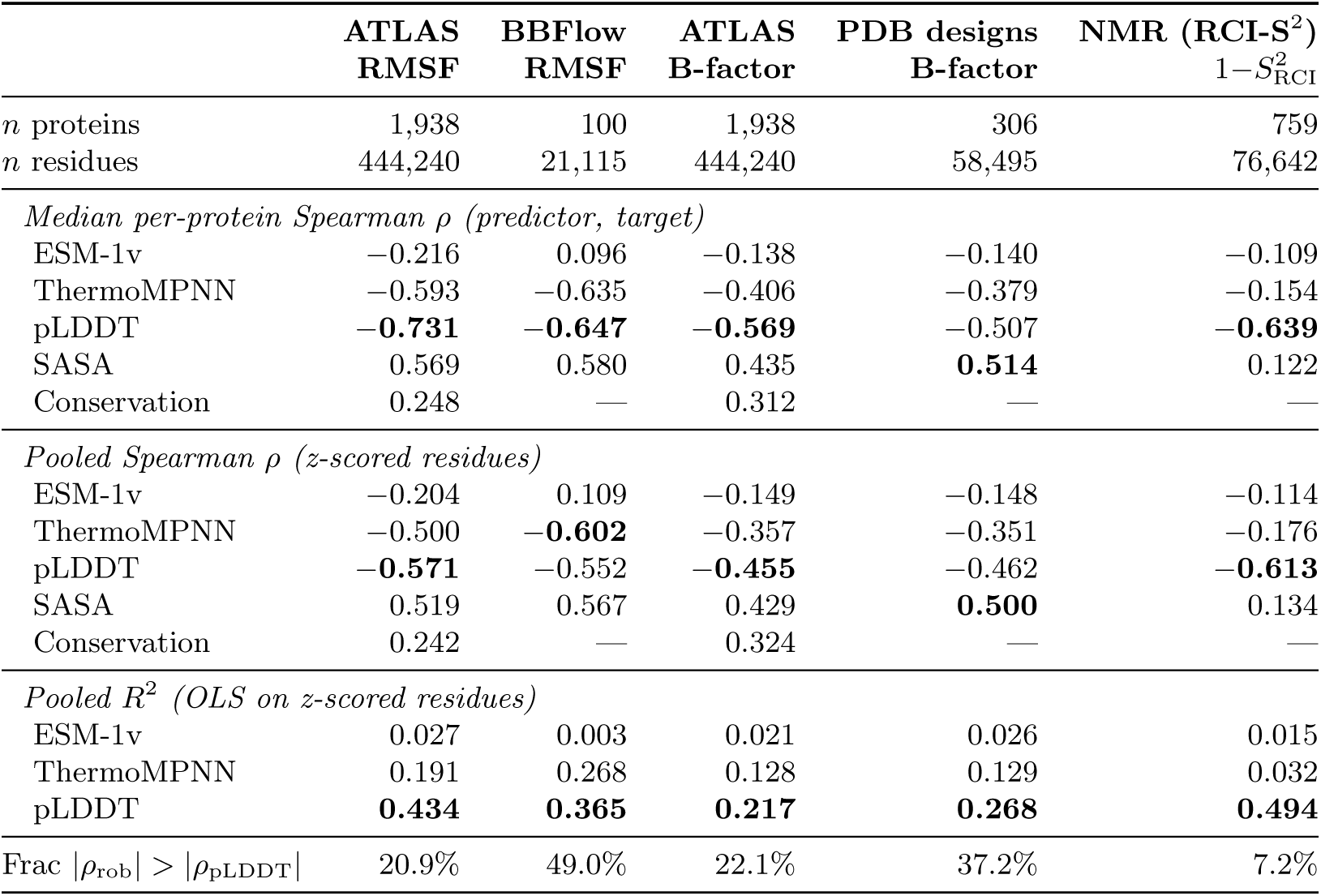
Bivariate correlations between per-residue predictors and dynamics targets across all dataset–target combinations. *Median per-protein ρ*: median across proteins of the within-protein Spearman rank correlation. *Pooled ρ*: Spearman correlation on all residues after within-protein z-scoring. *Pooled R*^2^: OLS on z-scored residues. Conservation scores from ConSurf Rate4Site (ATLAS only). Best value in each predictor comparison is shown in bold. All pooled correlations significant at *p <* 10^−10^. Partial correlations and incremental *R*^2^ are in Supplementary Table S1.

### 2.4 Statistical Analysis

For each protein, we computed the Spearman rank correlation *ρ* between the per-residue robustness index *R_i_* and the dynamics measure (RMSF*_i_*or *B_i_*) across positions *i* = 1*, …, L*. We report the median *ρ* across all proteins as the primary metric.

For pooled analysis, we computed per-residue z-scores within each protein, then combined all residues from all proteins. Within-protein z-scoring was necessary because proteins differ widely in their overall stability and dynamics scale: a highly stable protein may have uniformly large |ΔΔ*G*| and low RMSF, while a marginally stable protein may have the opposite. Without normalization, these protein-level offsets would dominate the pooled signal and obscure the within-protein residue-level relationship of interest.

We report both Spearman *ρ* and *R*^2^ (from OLS regression on z-scored values): Spearman *ρ* measures the strength of the monotonic association (robust to nonlinearity and outliers), while *R*^2^ quantifies the proportion of variance explained by a linear model. We also computed joint *R*^2^ from linear regression of the dynamics target on both robustness and a covariate, to assess incremental variance explained (Δ*R*^2^).

To isolate the robustness signal from confounders, we computed partial Spearman correlations in three setups: (i) *ρ*(robustness, dynamics | pLDDT), testing whether robustness carries information beyond structural confidence; (ii) *ρ*(robustness, dynamics | SASA), testing whether the signal survives controlling for burial; and (iii) *ρ*(robustness, dynamics | pLDDT + SASA), the most conservative control. All reported *p*-values are from two-sided tests. Because we report a modest number of correlations (*<* 50 across all target vs. covariate combinations) and all observed *p*-values are below 10^−6^, multiple-testing correction (e.g. Bonferroni) does not alter any conclusion.

We also stratified residues by secondary structure (*α*-helix H, *β*-sheet E, coil C; assigned using DSSP via *mdtraj*) and by burial class based on relative solvent accessibility (RSA): core (RSA *<* 20%), boundary (20-50%), and surface (*>* 50%), cutoffs that yield approximately equal-sized groups.

### 2.5 Multi-ΔΔ*G* Regression

To test whether the full per-residue mutational profile carries more information than any scalar summary, we constructed a 20-dimensional feature vector **x***_i_* = (ΔΔ*G_i,_*_A_, ΔΔ*G_i,_*_C_*, …,* ΔΔ*G_i,_*_Y_) for each residue *i*, where each component is the predicted ΔΔ*G* for substitution to a specific target amino acid (the wild-type entry is zero). We compared the following regression models for predicting the dynamics target (RMSF or B-factor):

1. **OLS on** std(ΔΔ*G*): baseline univariate model.
2. **Ridge on 20** ΔΔ*G* **features**: the full mutational profile with *L*_2_ regularization (*α* = 1.0).
3. **Ridge on 20** ΔΔ*G* **+ nonlinear features**: the 20 per-amino-acid values augmented with four nonlinear summary statistics: std(ΔΔ*G*), mean |ΔΔ*G*|, max |ΔΔ*G*|, and min(ΔΔ*G*), yielding 24 features.
4. Models with **pLDDT** added as an additional feature.

We evaluated all models in 5-fold protein-level cross-validation: we randomly assigned proteins (not residues) to folds, so the model must generalize to entirely unseen proteins. We z-scored features and target within each protein before pooling, matching the normalization used in the bivariate analysis (Section 3.1), so the regression captures within-protein residue-level signal rather than between-protein offsets. Ridge regularization prevents overfitting on the 20-25 dimensional feature space. We averaged per-amino-acid coefficients across folds and report them as standardized regression weights, as shown in Figure 3.

## 3 Results

### 3.1 Bivariate Correlations

Table 1 summarizes the main correlations between std(ΔΔ*G*) robustness and dynamics across all dataset-target combinations.

ThermoMPNN robustness shows a consistent negative correlation with RMSF: median per-protein *ρ* = −0.593 on ATLAS and −0.635 on BBFlow, the latter nearly matching pLDDT (−0.647). In pooled analysis, ThermoMPNN explains *R*^2^ = 0.191 (ATLAS RMSF, 444,240 residues) and *R*^2^ = 0.268 (BBFlow, 21,115 residues). ESM-1v was much weaker: median *ρ* = −0.216 on ATLAS (pooled *R*^2^ = 0.027, 7-fold lower than ThermoMPNN) and produced *wrong-sign* correlations on designed proteins (*ρ* = +0.096 on BBFlow), indicating that a sequence-only model cannot capture the robustness-dynamics coupling in proteins that lack an evolutionary sequence record. On the NMR dataset, ESM-1v likewise underperformed ThermoMPNN (Table 1). All subsequent analyses therefore use ThermoMPNN only.

B-factor correlations followed the same pattern at reduced magnitude, as expected given the noisier nature of crystallographic B-factors [Sun et al., 2019, Vander Meersche et al., 2025a].

Figure 1 shows the distribution of per-protein correlations. pLDDT (orange) concentrates at strongly negative *ρ* values, while robustness (blue) is broader and centered closer to zero; nearly all proteins lie below the diagonal, confirming that pLDDT is the stronger per-protein descriptor. The NMR dataset shows a markedly broader robustness distribution, with 31% of proteins exhibiting positive (wrong-sign) *ρ* (Section 3.6; Supplementary Figure S4). Figure 2 shows the pooled residue-level relationship after within-protein z-scoring; the raw scatter in Supplementary Figure S1 confirms that without z-scoring, between-protein variation dominates.

**Figure 1:**
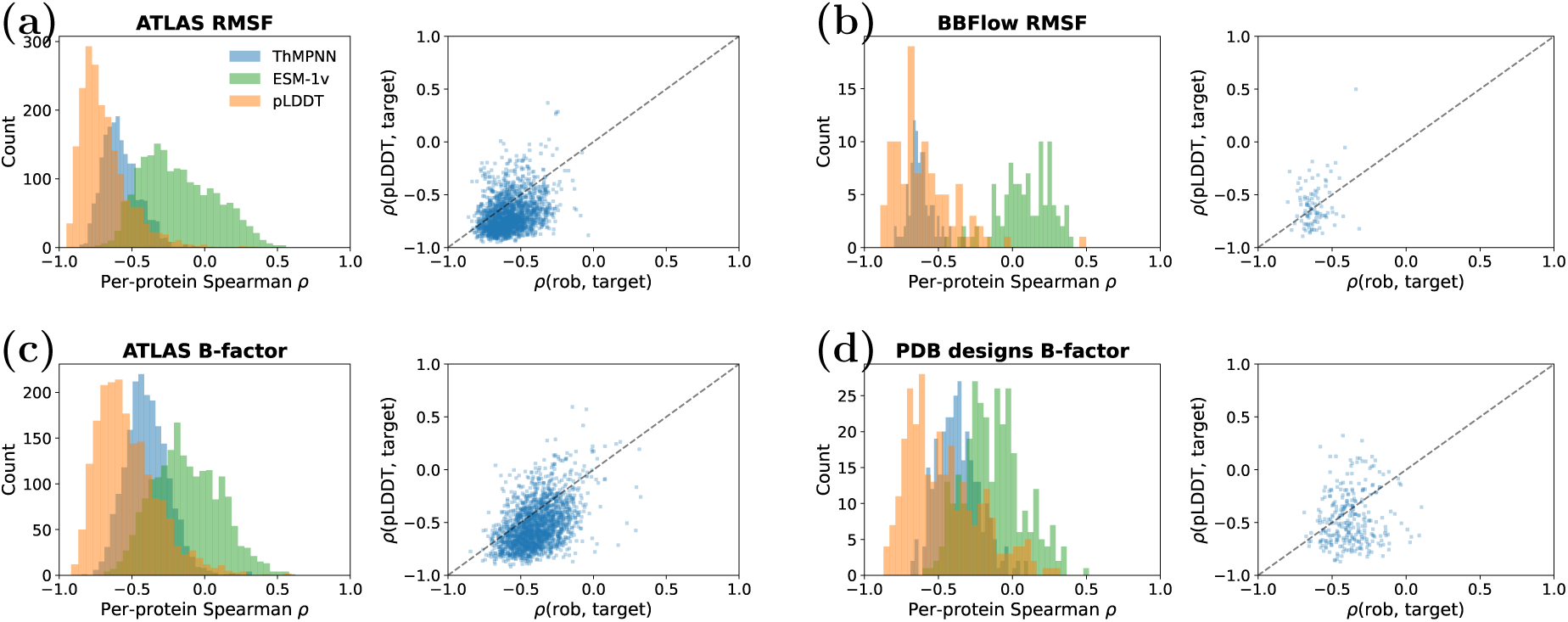
Distribution of within-protein Spearman *ρ* between predictor and dynamics target. Each panel shows one dataset-target pair. *Left:* histograms of per-protein *ρ* for ThermoMPNN robustness (blue), ESM-1v robustness (green, where available), and pLDDT (orange). *Right:* scatter of robustness *ρ* vs. pLDDT *ρ* per protein; points below the diagonal indicate pLDDT is the stronger descriptor. **(a)** ATLAS RMSF (*n* = 1,938), **(b)** BBFlow RMSF (*n* = 100), **(c)** ATLAS B-factor (*n* = 1,938), **(d)** PDB de novo designs B-factor (*n* = 306; pLDDT from ESMFold, *n* = 290). NMR results are in Supplementary Figure S4.

**Figure 2:**
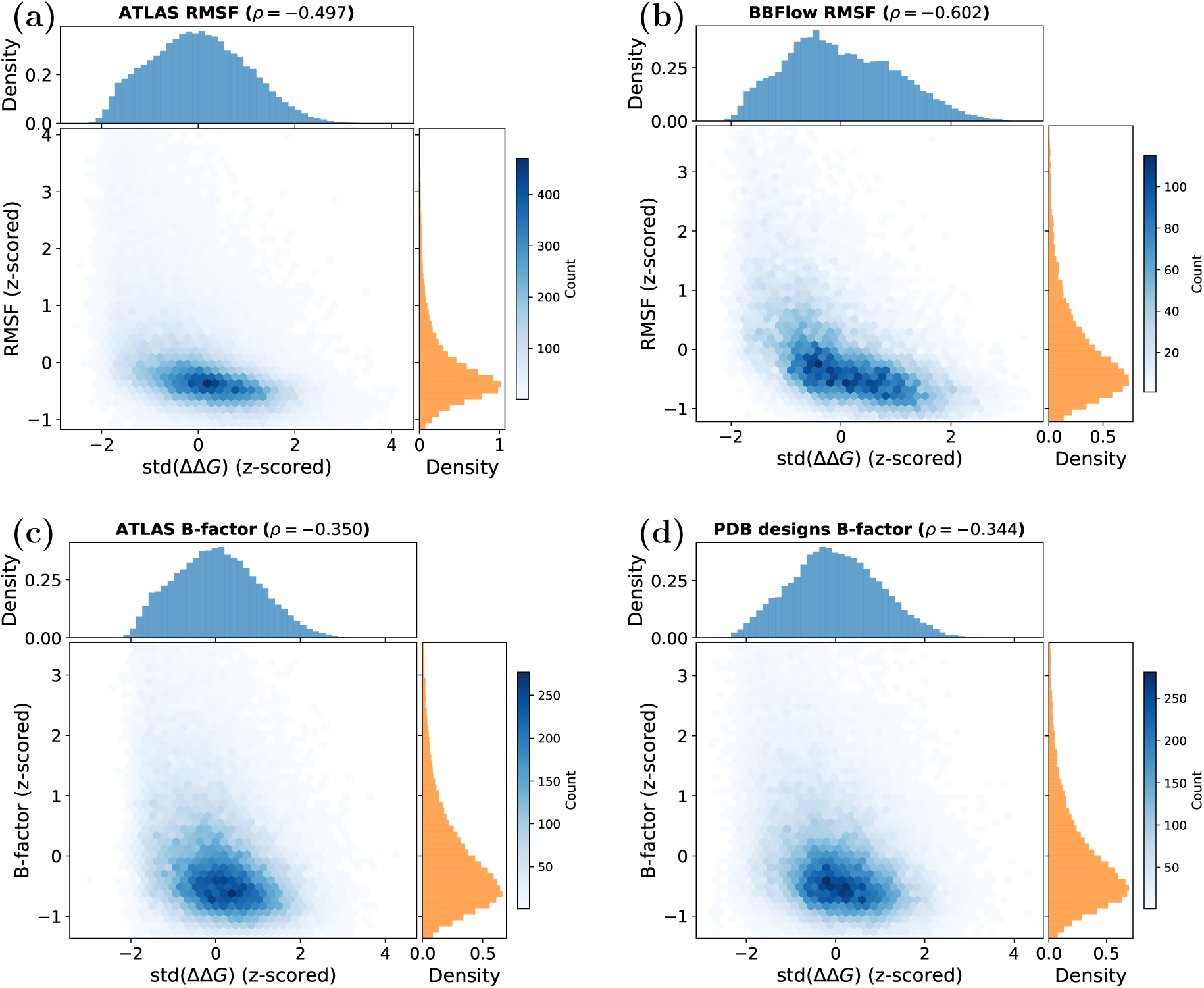
Pooled 2D density (hex-bin) of per-residue std(ΔΔ*G*) (z-scored within protein) vs. dynamics target (z-scored) for ThermoMPNN. Marginal histograms are shown on the top and right axes. The *y*-axis is clipped at the 1st and 99th percentiles to reduce heavy-tail distortion. **(a)** ATLAS RMSF, **(b)** BBFlow RMSF, **(c)** ATLAS B-factor, **(d)** PDB de novo designs B-factor. Pooled Spearman *ρ* and sample size are shown in each panel title. NMR results are in Supplementary Figure S5.

### 3.2 Partial Correlations and Incremental *R*^2^

pLDDT is the single strongest descriptor of dynamics in pooled analysis (ATLAS RMSF: *ρ* = −0.571, *R*^2^ = 0.434). However, adding ThermoMPNN std(ΔΔ*G*) to a joint model consistently increases *R*^2^ across all five dataset-target combinations, with the largest gain on BBFlow RMSF (Δ*R*^2^ = 0.104; Supplementary Table S1). On a per-protein basis, robustness beats pLDDT (|*ρ*_rob_| *>* |*ρ*_pLDDT_|) for 49% of BBFlow designed proteins vs. 21% of ATLAS natural proteins (RMSF target), and for 37% of PDB designs vs. 22% of ATLAS (B-factor target).

Partial correlations confirm that the robustness signal is not redundant with either pLDDT or SASA (Supplementary Table S1). After controlling for pLDDT, ThermoMPNN retains partial *ρ* between −0.09 (NMR) and −0.52 (BBFlow) across datasets. After controlling for SASA, partial *ρ* ranges from −0.12 (NMR) to −0.36 (BBFlow). The NMR partials are weaker, consistent with the overall weaker robustness signal on chemical-shift-derived dynamics (Section 3.6). On the MD-based datasets, the SASA partials are slightly weaker than the pLDDT partials, reflecting the collinearity between robustness and burial (both capture local packing density), while pLDDT is a model confidence score that does not directly probe the energy landscape, so robustness contributes more complementary information. All partial correlations are significant (*p <* 10^−6^).

### 3.3 Stratification by Secondary Structure and Burial

Table 2 shows pooled Spearman *ρ* stratified by secondary structure and burial class, with pLDDT values in parentheses. By secondary structure, robustness is strongest in *α*-helices (*ρ* = −0.40 on ATLAS RMSF, −0.49 on BBFlow) and coils (−0.45, −0.47), and weakest in *β*-sheets (−0.33, −0.45). By burial class, surface residues carry the strongest signal (*ρ* = −0.44 on ATLAS RMSF, −0.49 on BBFlow), while core residues show weak correlations (−0.23, −0.30), since they are uniformly rigid and mutationally sensitive, leaving little variance to explain. On BBFlow, the robustness-pLDDT gap narrows: in helices, robustness (−0.490) slightly exceeds pLDDT (−0.451), and in sheets they are nearly equal (−0.450 vs. −0.454). This pattern is consistent across all dataset-target combinations.

**Table 2:** Stratified pooled Spearman *ρ* between ThermoMPNN std(ΔΔ*G*) and dynamics target, grouped by secondary structure (DSSP classification: *α*-helix H, *β*-sheet E, coil C) and by burial class based on relative solvent accessibility (RSA): core (RSA *<* 20%), boundary (20–50%), surface (*>* 50%). pLDDT values shown in parentheses where available. Best |*ρ*| within each column (across structural classes) is shown in bold.

|  | ATLAS<br>RMSF | BBFlow<br>RMSF | ATLAS<br>B-factor | PDB designs<br>B-factor | NMR (RCI-S<sup>2</sup>)<br>1-S<sup>2</sup><sub>RCI</sub> |
| --- | --- | --- | --- | --- | --- |
| *Secondary structure* |  |  |  |  |  |
| α-helix | -0.402 (-0.469) | **-0.490 (-0.451)** | -0.287 (-0.383) | **-0.290 (-0.363)** | -0.088 (-0.562) |
| β-sheet | -0.333 (-0.439) | -0.450 (-0.454) | -0.248 (-0.352) | -0.177 (-0.355) | -0.114 (-0.533) |
| Coil | **-0.448 (-0.586)** | -0.466 (-0.472) | **-0.296 (-0.459)** | -0.289 (-0.459) | **-0.170 (-0.659)** |
| *Burial (RSA cutoffs)* |  |  |  |  |  |
| Core | -0.230 (-0.407) | -0.301 (-0.441) | -0.110 (-0.304) | -0.111 (-0.318) | -0.077 (-0.576) |
| Boundary | -0.338 (-0.487) | -0.380 (-0.509) | -0.206 (-0.378) | -0.212 (-0.374) | -0.130 (-0.600) |
| Surface | **-0.444 (-0.590)** | **-0.488 (-0.590)** | **-0.286 (-0.438)** | **-0.298 (-0.436)** | **-0.213 (-0.646)** |

The BBFlow dataset lacks crystal structures, so the designed-protein MD results cannot be directly compared with the ATLAS B-factor analysis. To close this gap, we applied the same analysis to 306 de novo designed proteins crystallized in the PDB (Section 2.3), using B-factor as the dynamics target, enabling a direct natural-vs.-designed comparison with the same experimental measure. The pooled effect sizes are nearly identical: *ρ* = −0.351, *R*^2^ = 0.129 on PDB designs vs. *ρ* = −0.357, *R*^2^ = 0.128 on ATLAS (Table 1). Partial correlations after controlling for pLDDT (−0.254 vs. −0.209) and SASA (−0.181 vs. −0.175; Supplementary Table S1) are likewise comparable. Stratification mirrors the ATLAS pattern: helices (*ρ* = −0.290) and coils (−0.289) carry the signal while *β*-sheets are weaker (−0.177), and surface residues drive the strongest correlations (−0.298; Table 2). This consistency across natural and designed proteins, and across computational and experimental dynamics measures, supports a biophysical origin rather than an artifact of evolutionary biases.

### 3.4 Alternative Robustness Measures and Multi-ΔΔ*G* Regression

We first compare scalar summaries of the per-residue ΔΔ*G* distribution (Table 3, top panel), then test whether the full 20-dimensional ΔΔ*G* profile **x***_i_* = (ΔΔ*G_i,_*_A_*, …,* ΔΔ*G_i,_*_Y_) carries additional predictive information (bottom panel; 5-fold protein-level CV, ThermoMPNN scorer).

**Table 3:**
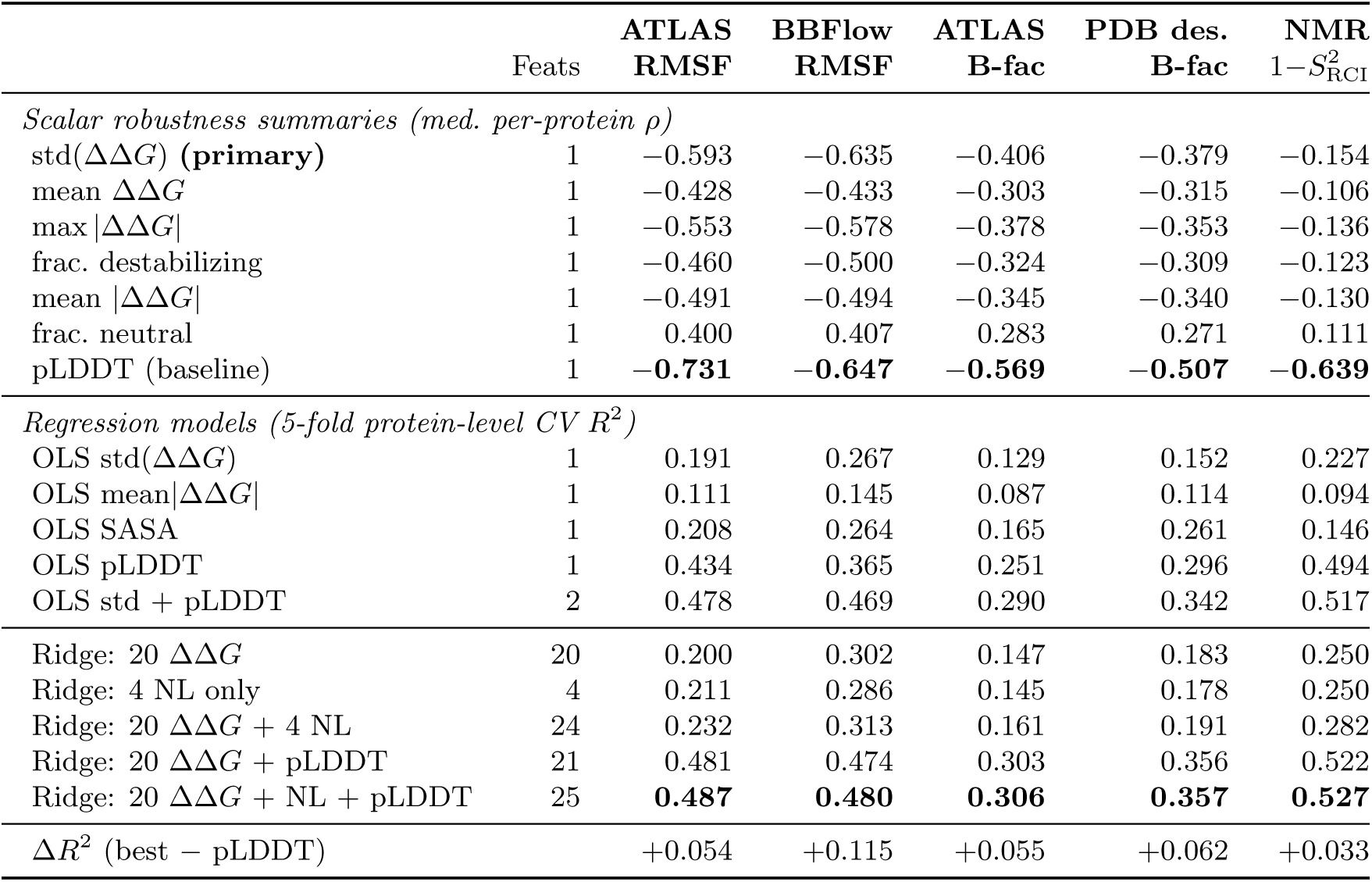
Comparison of robustness summary statistics and multi-ΔΔ*G* regression models (ThermoMPNN scorer). *Top panel* : median per-protein Spearman *ρ* for scalar robustness summaries. *Bottom panel* : 5-fold protein-level cross-validated *R*^2^ for regression models predicting dynamics from ΔΔ*G* features; proteins are held out as entire units so the model must generalize to unseen proteins. “Feats” = number of input features. Best value in each column section is shown in bold.

The standard deviation std(ΔΔ*G*) consistently outperforms all scalar alternatives (Table 3, top panel): on ATLAS RMSF, median *ρ* = −0.593 vs. −0.553 for max |ΔΔ*G*|, −0.491 for mean |ΔΔ*G*|, −0.460 for frac. destabilizing, and −0.400 for frac. neutral. On BBFlow, std(ΔΔ*G*) (−0.635) nearly matches pLDDT (−0.647), while all other robustness summaries are weaker than −0.58. The mean |ΔΔ*G*| is weaker because it does not distinguish positions where all mutations are moderately destabilizing from positions with a mix of neutral and catastrophic mutations; the standard deviation captures this heterogeneity.

Table 3 (bottom panel) and Figure 3 compare regression models in 5-fold protein-level cross-validation. On ATLAS RMSF, the 20-ΔΔ*G* Ridge model achieves CV *R*^2^ = 0.200, slightly above OLS on std(ΔΔ*G*) alone (*R*^2^ = 0.191). On BBFlow RMSF, the gain is larger: 20-ΔΔ*G* (*R*^2^ = 0.302), and the best combined model (20-ΔΔ*G* + nonlinear + pLDDT) reaches *R*^2^ = 0.480, a Δ*R*^2^ = +0.115 over pLDDT alone (*R*^2^ = 0.365). This gain is 2-fold larger than on ATLAS (+0.054), suggesting that the ΔΔ*G* profile captures dynamical information that pLDDT misses on designed proteins. On PDB designs (B-factor), the 20-ΔΔ*G* + nonlinear model (*R*^2^ = 0.191) outperforms the scalar std(ΔΔ*G*) (*R*^2^ = 0.152), and the best combined model (*R*^2^ = 0.357) exceeds pLDDT alone (*R*^2^ = 0.296) with Δ*R*^2^ = +0.062. B-factor results mirror RMSF with reduced *R*^2^ values and identical model ranking (Table 3), strengthening confidence that the ΔΔ*G* profile captures genuine dynamical information rather than MD force-field artifacts.

**Figure 3:**
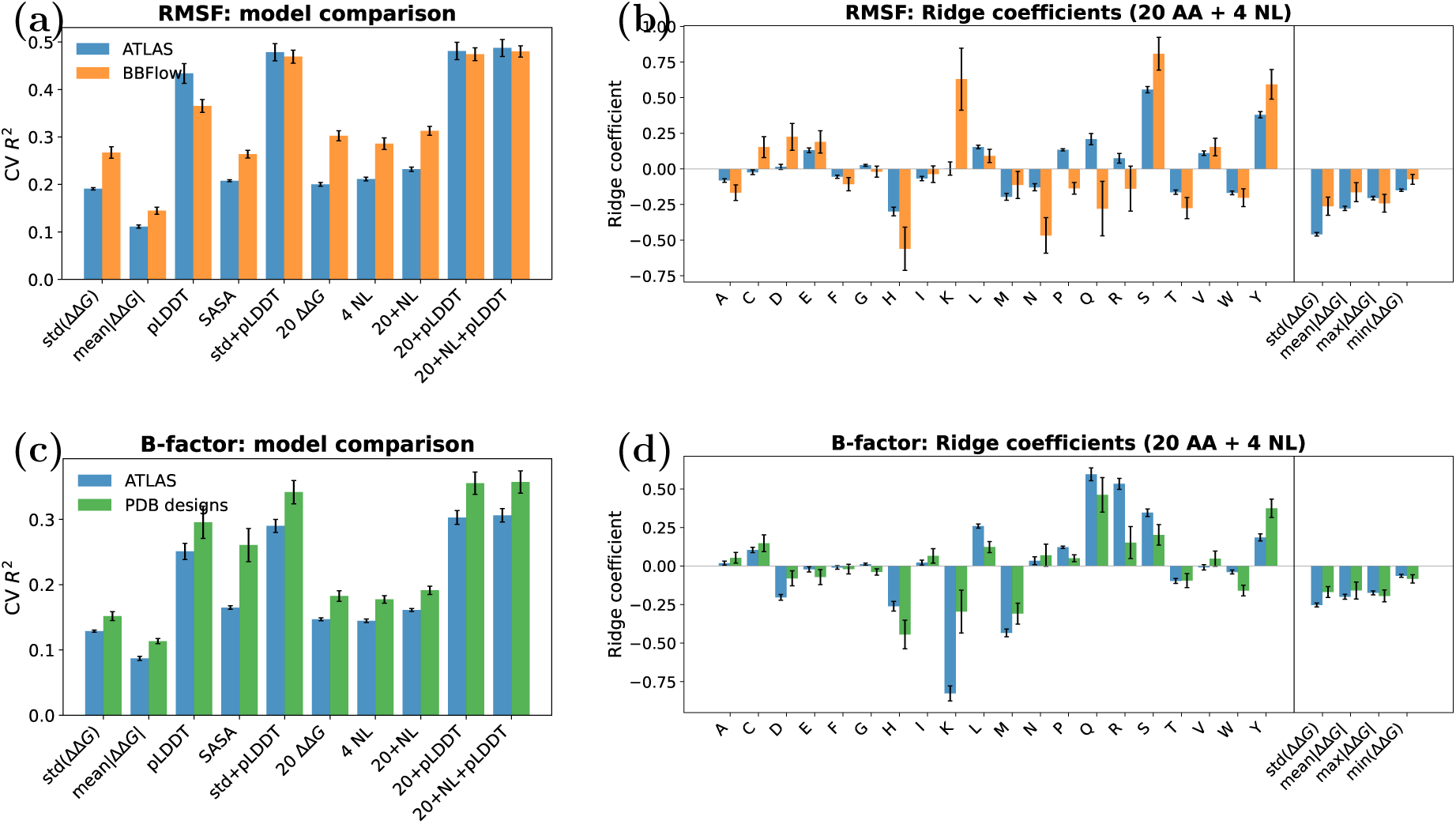
Multi-ΔΔ*G* regression results (5-fold protein-level CV, ThermoMPNN). Two rows: RMSF and B-factor. *Left column:* cross-validated *R*^2^ for scalar baselines and multi-feature models; error bars show standard deviation across folds. *Right column:* Ridge coefficients for the 24-feature model (20 per-amino-acid ΔΔ*G* values + 4 nonlinear summaries, separated by vertical line); error bars show ±2 SE. Datasets: ATLAS (blue), BBFlow (orange), PDB designs (green). Positive coefficients: costly mutation to that amino acid associated with flexibility; negative: rigidity. NMR results are in Supplementary Figure S6.

Figure 3 shows Ridge coefficients from the 24-feature model (20 per-amino-acid ΔΔ*G* values + 4 nonlinear summary statistics), so that per-AA and summary-statistic contributions are on the same scale. For RMSF, the strongest positive per-AA coefficients correspond to S (serine, +0.81 on BBFlow, +0.56 on ATLAS) and Y (tyrosine, +0.59, +0.38): positions where mutation *to* these amino acids is costly tend to be more dynamic. The strongest negative per-AA coefficients are H (histidine, −0.56 on BBFlow, −0.30 on ATLAS) and N (asparagine, −0.47 on BBFlow). Among the nonlinear summaries, std(ΔΔ*G*) carries the largest coefficient on ATLAS RMSF (−0.46). For B-factor, K (lysine) emerges as the single largest per-AA signal (−0.83 on ATLAS), while Q (glutamine, +0.60) and R (arginine, +0.53) are the strongest positive. The ATLAS and BBFlow coefficient profiles are broadly concordant for RMSF, and ATLAS and PDB designs agree for B-factor, providing cross-dataset validation. On the NMR 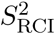 dataset (Supplementary Figure S6), std(ΔΔ*G*) dominates (−0.63), with per-AA coefficients (H −0.46, S +0.40, Y +0.39) playing a secondary role; the model achieves CV *R*^2^ = 0.282, confirming that amino-acid-specific mutational patterns capture NMR dynamics information that scalar summaries miss.

### 3.5 Case Studies: Proteins Where Robustness Outperforms pLDDT

While pLDDT is the strongest single predictor of dynamics in pooled analysis (Section 3.3), robustness outperforms pLDDT on a substantial minority of individual proteins: |*ρ*_rob_| *>* |*ρ*_pLDDT_| for 21% of ATLAS proteins (RMSF target) and 49% of BBFlow designed proteins. To illustrate the per-residue robustness signal on proteins where it adds the most value over pLDDT, we selected three ATLAS proteins where robustness substantially outperforms pLDDT on both RMSF and B-factor targets.

The Zika virus capsid protein (PDB 5YGH, chain A, 76 residues; Figure 4) is a structural component of the viral particle that mediates RNA binding and virus assembly. The capsid protein undergoes conformational changes during viral maturation, and its dynamics are relevant for understanding viral assembly mechanisms and potential drug targets. On this protein, robustness outperforms pLDDT: the robustness index achieves *ρ*_RMSF_ = −0.72 and *ρ*_B-factor_ = −0.67, while pLDDT achieves only *ρ*_RMSF_ = −0.29 and has a near-zero *positive* (wrong-sign) correlation with B-factor (*ρ* = +0.04). This means that pLDDT’s per-residue confidence scores carry essentially no information about local flexibility in this protein, whereas the mutational landscape correctly identifies which residues are dynamic. The robustness-colored structure, displayed in Figure 4c, shows clear correspondence with the RMSF and B-factor panels (d, e), with flexible loop regions in red and the helical core in blue, while the pLDDT-colored structure (b) appears nearly uniformly blue (high confidence throughout).

**Figure 4:**
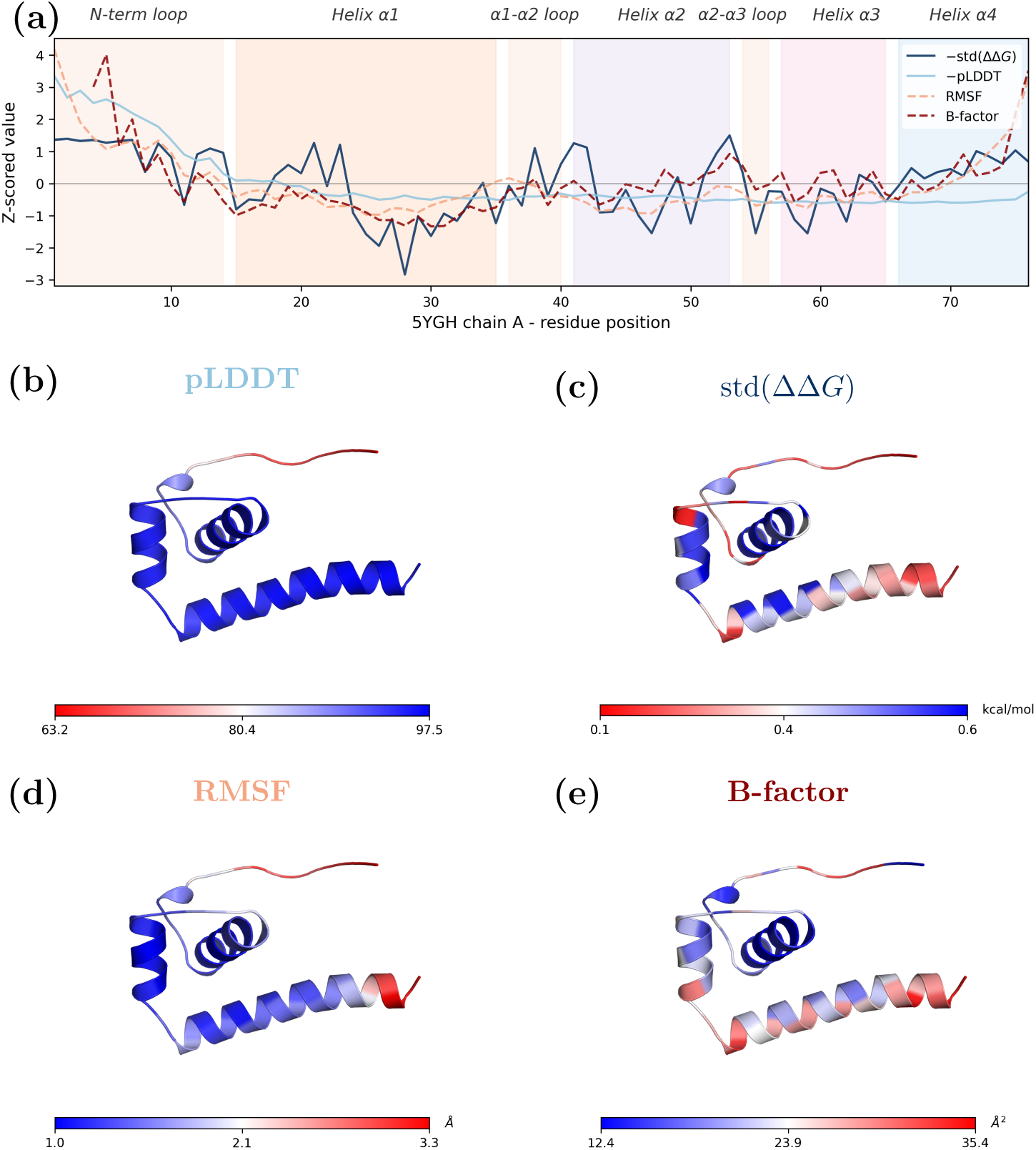
Per-residue robustness vs. dynamics on the Zika virus capsid protein (PDB 5YGH, chain A, 76 residues). **(a)** Z-scored per-residue profiles. **(b)**-**(e)** 3D structure colored by each metric: predictors on top (b, c), targets on bottom (d, e). Blue = rigid, red = flexible. pLDDT has a near-zero *positive* (wrong-sign) correlation with B-factor (*ρ* = +0.04), indicating that its per-residue confidence scores carry no useful information about flexibility on this protein, while robustness achieves *ρ*_RMSF_ = −0.72 and *ρ*_B-factor_ = −0.67.

The robustness trace, shown in Figure 4a, exhibits a ∼3.6-residue oscillation within helical segments, reflecting the alternation between buried and solvent-exposed side-chain positions on opposite faces of the *α*-helix. This face-dependent burial asymmetry is captured by std(ΔΔ*G*) because it is sensitive to local side-chain packing, whereas backbone dynamics measures (RMSF, B-factor) average over the helix as a relatively rigid body and show no such oscillation. This observation is consistent with the partial correlation analysis (Supplementary Table S1), where controlling for SASA reduces but does not eliminate the robustness-dynamics signal.

The vaccinia virus immunomodulator A46 N-terminal domain (PDB 5EZU, chain A, 89 residues; Supplementary Figure S2) sequesters host TIR-domain proteins to evade innate immune signaling. This *β*-sheet-rich domain also harbors a lipid-binding site for myristic acid. Robustness achieves *ρ*_RMSF_ = −0.75 and *ρ*_B-factor_ = −0.44, substantially outperforming pLDDT (*ρ*_RMSF_ = −0.30, *ρ*_B-factor_ = −0.26). The helix-face oscillation is less prominent in this predominantly *β*-sheet structure, as expected.

The L27 domain of Discs Large 1 (DLG1) (PDB 4RP5, chain A, 98 residues; Supplementary Figure S3) is a self-assembly module from the *Drosophila* tumor suppressor scaffolding protein DLG1, which mediates protein-protein interactions critical for epithelial cell polarity and growth control. Robustness achieves *ρ*_RMSF_ = −0.78 and *ρ*_B-factor_ = −0.33, while pLDDT reaches only *ρ*_RMSF_ = −0.26 and has essentially zero correlation with B-factor (*ρ* = +0.01).

These case studies demonstrate that on proteins where pLDDT fails to capture local flexibility, the mutational robustness index provides a complementary and often superior signal, mapping physically meaningful dynamical patterns onto the three-dimensional structure.

### 3.6 Validation on NMR-derived Dynamics 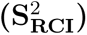

To test the robustness-dynamics relationship against an independent experimental modality, we analyzed the 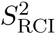 dataset of Gavalda-Garcia et al. [2025], which provides NMR chemical-shift-derived order parameters for 759 proteins (76,642 residues). We used 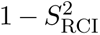 as the dynamics target so that high values indicate flexibility, matching the RMSF and B-factor convention.

The scalar robustness index std(ΔΔ*G*) shows a weak pooled correlation with 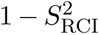 (*ρ* = −0.176, *R*^2^ = 0.032), substantially below the ATLAS RMSF and B-factor results (Table 1). In the CV regression framework, the 20-ΔΔ*G* + 4 nonlinear Ridge model achieves CV *R*^2^ = 0.282, and the best combined model (with pLDDT) reaches *R*^2^ = 0.527 (Table 3). pLDDT alone achieves *ρ* = −0.613 (*R*^2^ = 0.494) on this dataset, stronger than on ATLAS RMSF, likely because 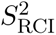 captures order/disorder classification (where pLDDT excels) rather than the fine-grained dynamical gradations within ordered regions that MD-RMSF resolves. The ΔΔ*G* profile still adds Δ*R*^2^ = +0.033 over pLDDT alone.

The gap between the pooled *R*^2^ (0.032) and the CV *R*^2^ for OLS std(ΔΔ*G*) (0.227) is unusually large on this dataset. The source is visible in Supplementary Figure S4: the median per-protein *ρ* is only −0.154, with 31% of proteins showing *positive* (wrong-sign) correlations, compared to fewer than 1% on ATLAS and BBFlow. In the pooled analysis, residues from proteins with opposite-sign correlations partially cancel, depressing the aggregate *R*^2^; in the CV framework, protein-level averaging preserves the signal from the majority of proteins where the expected negative correlation holds.

The weak scalar but meaningful 20-dimensional signal suggests that while the overall variance of the mutational landscape is only weakly coupled to chemical-shift-derived order parameters, the *shape* of the landscape (which amino acids are costly vs. tolerated) encodes structural constraint information that aligns with NMR dynamics, supporting the generality of the multi-ΔΔ*G* approach across experimental modalities.

## 4 Discussion

In this study, we examined the hypothesis presented in Tokuriki and Tawfik [2009a], that local mutational robustness quantitatively predicts local protein dynamics at the single residue level. Our results confirm a consistent negative correlation between robustness and flexibility, with effect size depending on the scorer, the dynamics target, and how the mutational profile is summarized.

We have shown that ThermoMPNN substantially outperforms ESM-1v: a 3-fold difference in median |*ρ*| on natural proteins and a sign reversal on designed proteins (Table 1). A sequence-only model has no access to backbone geometry, the primary determinant of both stability and dynamics, and de novo designs lack the evolutionary sequence record that ESM-1v implicitly relies on. Studies using sequence-only ΔΔ*G* predictors to characterize designed proteins should therefore be interpreted with caution. However, newer sequence-only predictors such as SPURS [Li and Luo, 2025] report even higher accuracy than ThermoMPNN on experimental ΔΔ*G* benchmarks; whether their improved accuracy translates into stronger robustness-dynamics correlations, particularly on designed proteins that lack evolutionary sequence records, remains to be tested.

pLDDT is the strongest single descriptor of dynamics (*R*^2^ = 0.434 for ATLAS RMSF), but robustness carries complementary information. The incremental Δ*R*^2^ from adding robustness to pLDDT is 2-fold larger on designed proteins than on natural ones (Supplementary Table S1), and partial correlations after controlling for pLDDT remain substantial on all datasets. Per-protein analysis shows that the pLDDT-robustness gap narrows on more flexible proteins, consistent with pLDDT capturing order/disorder classification but not gradations of flexibility within dynamic regions [Vander Meersche et al., 2025a, Gavalda-Garcia et al., 2025]; case studies on individual proteins where robustness substantially outperforms pLDDT confirm this pattern (Section 3.5). Robustness and SASA overlap more, since both reflect local packing density, while pLDDT does not directly probe the energy landscape. The 20-ΔΔ*G* Ridge model outperforms the best scalar summary on all datasets (Table 3), confirming that the identity of the target amino acid matters, and the concordance of Ridge coefficients across independent datasets supports a biophysical basis for the learned weights. The pooled effect sizes on natural and designed proteins are nearly identical when using the same experimental dynamics measure, and the signal survives controlling for pLDDT and SASA (Supplementary Table S1), supporting a structural origin rather than an evolutionary one.

Supervised tools such as PEGASUS [Vander Meersche et al., 2025b] achieve strong per-residue RMSF predictions (Pearson *r* ∼ 0.75) by training protein-language-model embeddings directly on MD-derived labels. Our robustness index is not trained on dynamics data at all, so the correlation with RMSF (median |*ρ*| ∼ 0.6) reveals a biophysical relationship rather than a learned association. A high std(ΔΔ*G*) directly indicates that a residue sits in a tightly packed environment sensitive to sidechain perturbation, providing mechanistic insight that black-box regressors do not. The two approaches are complementary: PEGASUS may achieve higher absolute correlation with MD targets, while robustness explains *why* a residue is dynamically constrained.

The per-residue std(ΔΔ*G*) quantifies the *curvature* of the stability landscape around the wild-type sequence: a high value indicates a rugged local landscape where substitutions have heterogeneous effects, while a low value indicates a flat landscape where most substitutions are tolerated. The correlation with dynamics shows that landscape ruggedness tracks local structural constraint: tightly packed, rigid residues sit in steep-sided stability wells, while flexible residues occupy flatter regions. This extends the protein-level robustness-evolvability framework of Bloom et al. [2006] and Wagner [2008] to the residue level: positions with flat local landscapes are both mutationally accessible and dynamically mobile, jointly facilitating sequence-space exploration during evolution. That designed proteins show the same pattern indicates a structural property of the fitness landscape itself, not a consequence of evolutionary optimization. The consistent burial-class gradient across all datasets likely reflects the low variance of both robustness and dynamics among uniformly rigid, mutationally sensitive core residues, while the weaker signal in *β*-sheets compared to helices and coils may arise from hydrogen-bond networks that constrain both mutational sensitivity and dynamical variation.

Our framework is agnostic to the choice of ΔΔ*G* predictor: the robustness index and the multi-ΔΔ*G* regression pipeline treat the scorer as a modular component, so improvements in ΔΔ*G* predictors are expected to translate into stronger robustness-dynamics correlations without methodological changes.

Robustness and conservation both reflect structural constraints, but they capture different quantities. Conservation records the *realized* evolutionary trajectory, shaped by selection, drift, and functional pressures beyond stability; robustness measures the *shape of the local energy landscape*, a biophysical property of the current structure independent of evolutionary history [Bloom and Arnold, 2009, Tokuriki and Tawfik, 2009b]. Active-site residues, for example, are often highly conserved due to functional selection yet thermodynamically tolerant of substitution because they sit in structurally relaxed environments.

Several lines of evidence argue that the robustness-dynamics signal is biophysical rather than a proxy for conservation. Robustness predicts dynamics equally well on designed proteins (BBFlow, PDB designs) that have no homologs and no evolutionary history, yet show comparable correlation magnitudes and partial correlations (Table 1; Supplementary Table S1). ESM-1v is itself a learned representation of evolutionary conservation [Meier et al., 2021], yet ThermoMPNN, which relies solely on three-dimensional structure, outperforms it 3-fold in median |*ρ*| on natural proteins and avoids the sign reversal on designed proteins; if conservation drove the link, an evolution-aware model should perform better, not worse. Moreover, the partial correlations controlling for pLDDT and SASA already remove much of the structural constraint signal that conservation shares with both robustness and dynamics, yet robustness remains a significant independent predictor.

We tested this directly by including per-residue ConSurf Rate4Site conservation scores [Ashkenazy et al., 2016] as an additional covariate in the ATLAS analyses. Conservation itself correlates moderately with dynamics (Table 1), but the partial correlation of robustness with dynamics after controlling for conservation remains substantial, and robustness explains additional variance (Δ*R*^2^) beyond a conservation-only model (Supplementary Table S1), confirming that the robustness signal is not merely a proxy for evolutionary constraint.

Our analysis is entirely within-protein: all features are z-scored per protein, so the native stability Δ*G* plays no role. This is appropriate for testing whether local mutational sensitivity predicts local dynamics [Tokuriki and Tawfik, 2009a], but does not address the protein-level prediction that more stable proteins should be globally more robust [Bloom et al., 2006, Wagner, 2008]. A between-protein analysis could be enabled by absolute Δ*G* predictors such as IFUM [Lee et al., 2026].

Our robustness index uses *predicted* ΔΔ*G* values, not experimental measurements, so the analysis partly tests how ThermoMPNN’s learned structural priors align with dynamical observables. Validation against experimental ΔΔ*G* data (e.g. MegaScale deep mutational scanning) would confirm that the signal reflects genuine thermodynamic sensitivity. Our primary dynamics measure (MD-RMSF) is itself computational; the B-factor analysis provides experimental grounding, but B-factors are noisy due to crystal-packing effects and refinement artifacts [Sun et al., 2019, Vander Meersche et al., 2025a]. The PDB de novo designs dataset is heterogeneous, spanning multiple design methods and assembled via keyword-based PDB search; some false positives may remain despite our filtering.

A particularly promising extension is to NMR relaxation data from the NMR-APP database [Müntener et al., 2026]. Crystallographic B-factors and MD-RMSF both capture small-amplitude fluctuations around a single equilibrium structure, whereas NMR relaxation rates (*R*_1_, *R*_2_, heteronuclear NOE) report on motions across a wide range of timescales: from picosecond bond vibrations to millisecond conformational exchange, and can detect transitions between distinct conformational substates on the energy landscape. Because robustness quantifies the local *curvature* of the stability landscape, it may be especially well suited to predict slow exchange motions that reflect barrier crossings between alternative conformations, a regime where pLDDT, which scores a single static fold, is expected to perform poorly [Vander Meersche et al., 2025a]. Testing this prediction would clarify whether the robustness-dynamics link extends beyond harmonic fluctuations to the large-scale conformational dynamics most relevant for function and allostery. Other promising directions include nonlinear models for the 20 ΔΔ*G* features, comparison against experimental ΔΔ*G* (e.g. the MegaScale dataset [Tsuboyama et al., 2023]), and application of robustness as a scoring function for conformational protein design.

## Data and Code Availability

All analysis code is available at https://github.com/orzuk/robustness-dynamics. ATLAS data are available from https://www.dsimb.inserm.fr/ATLAS/ [Vander Meersche et al., 2024]. BBFlow designed protein structures and MD trajectories are from Wolf et al. [2025]. PDB de novo design structures were retrieved from the RCSB PDB (https://www.rcsb.org/); the list of 306 PDB IDs used (after excluding 12 natural-protein contaminants) is provided in the code repository. 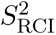 data are from the Zenodo repository of Gavalda-Garcia et al. [2025].

## Funding

This work was partly supported by the Israel Science Foundation (grant 2392/22).

## Acknowledgments

We thank Meira Barron for discussions on mutational robustness in the context of protein design, which motivated the investigation of robustness as a predictor of protein dynamics. We also thank Sarel Fleishman for suggesting the investigation of protein dynamics measures in the context of mutational robustness, and Barak Yariv and Nir Ben Tal for providing per-residue ConSurf conservation scores.

## A Supplementary Tables

**Table S1:**
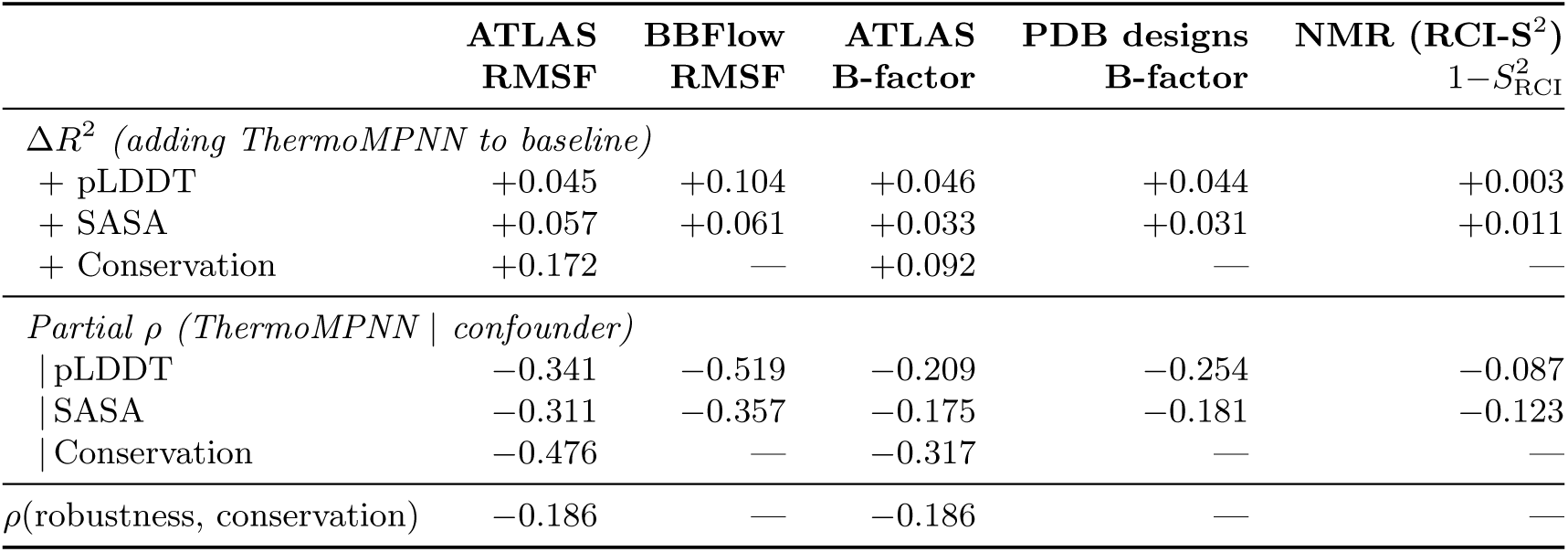
Partial correlations and incremental variance explained. Δ*R*^2^: increase in *R*^2^ when adding ThermoMPNN std(ΔΔ*G*) to the baseline predictor. *Partial ρ*: Spearman correlation of robustness vs. target after controlling for the indicated confounder. Conservation scores from ConSurf Rate4Site (ATLAS only). All partial correlations significant at *p <* 10^−6^.

## B Supplementary Figures

**Figure S1:**
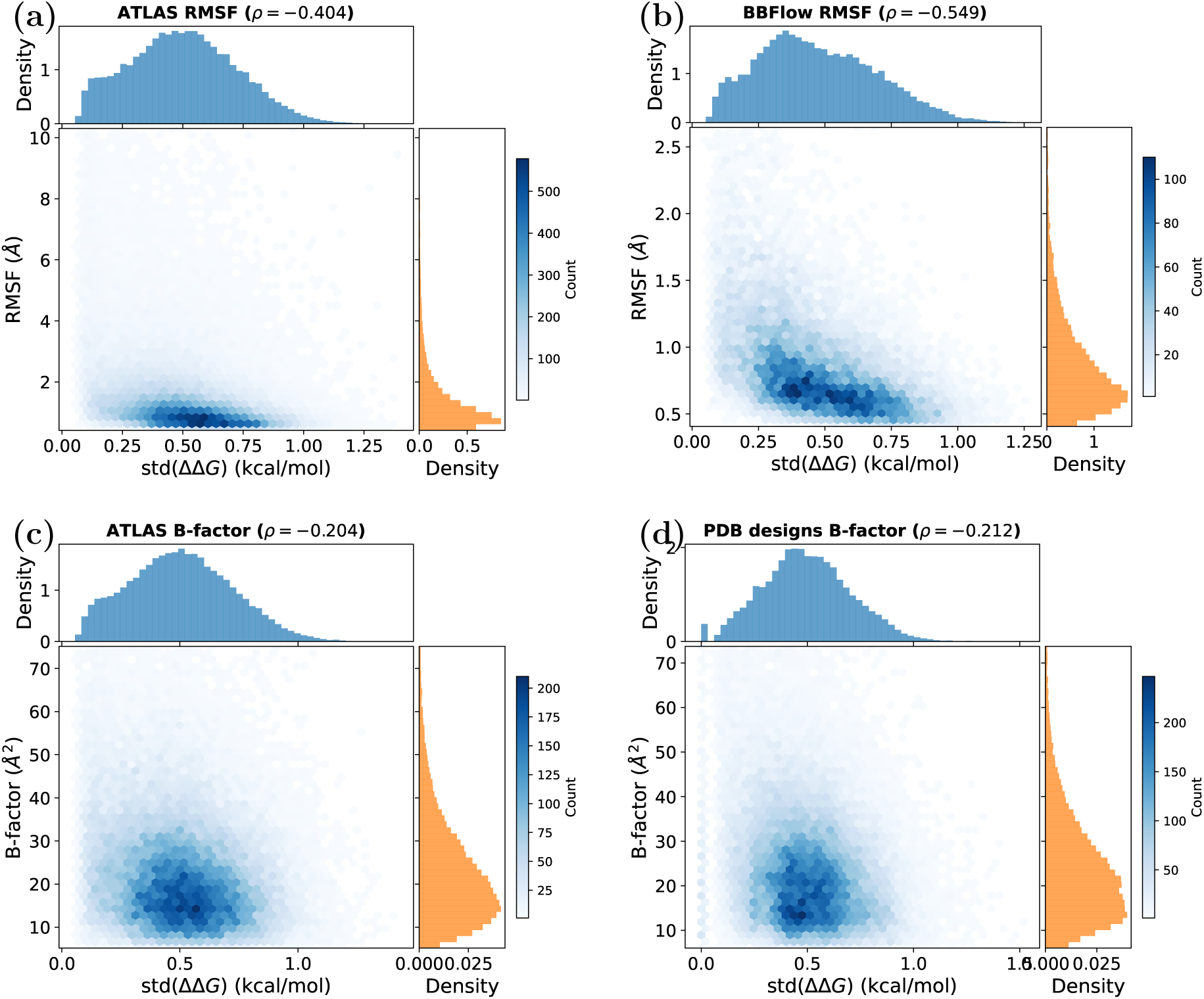
Pooled per-residue scatter of std(ΔΔ*G*) (kcal/mol) vs. dynamics target in raw (unnormalized) units. The data are dominated by between-protein variation: proteins with high overall RMSF or B-factor form distinct vertical bands. The raw Spearman *ρ* and sample size are shown in each panel title. All main-text analyses use within-protein z-scoring to remove protein-level offsets and isolate the residue-level robustness-dynamics relationship (Figure 2). **(a)** ATLAS RMSF, **(b)** BBFlow RMSF, **(c)** ATLAS B-factor, **(d)** PDB designs B-factor.

**Figure S2:**
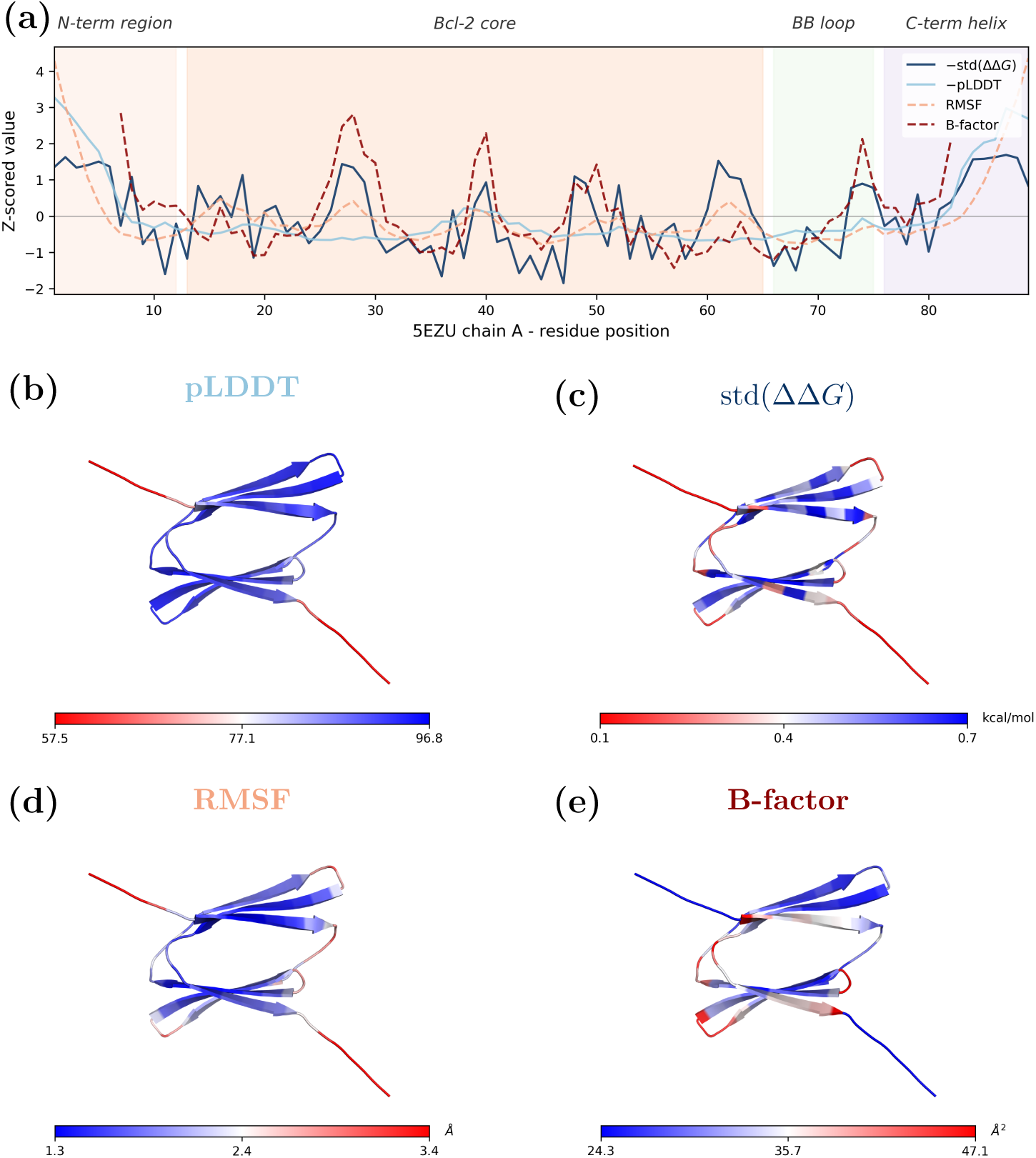
Per-residue robustness vs. dynamics on the vaccinia virus immunomodulator A46 N-terminal domain (PDB 5EZU, chain A, 89 residues). Layout as in Figure 4. Although the pLDDT-colored structure (b) visually resembles the B-factor pattern (e) in the structured core, the high-RMSF N-terminal coil (red in d) is missed by pLDDT, driving the overall correlation gap (*ρ*_rob,RMSF_ = −0.75 vs. *ρ*_pLDDT,RMSF_ = −0.30). The robustness panel (c) correctly identifies this flexible terminus.

**Figure S3:**
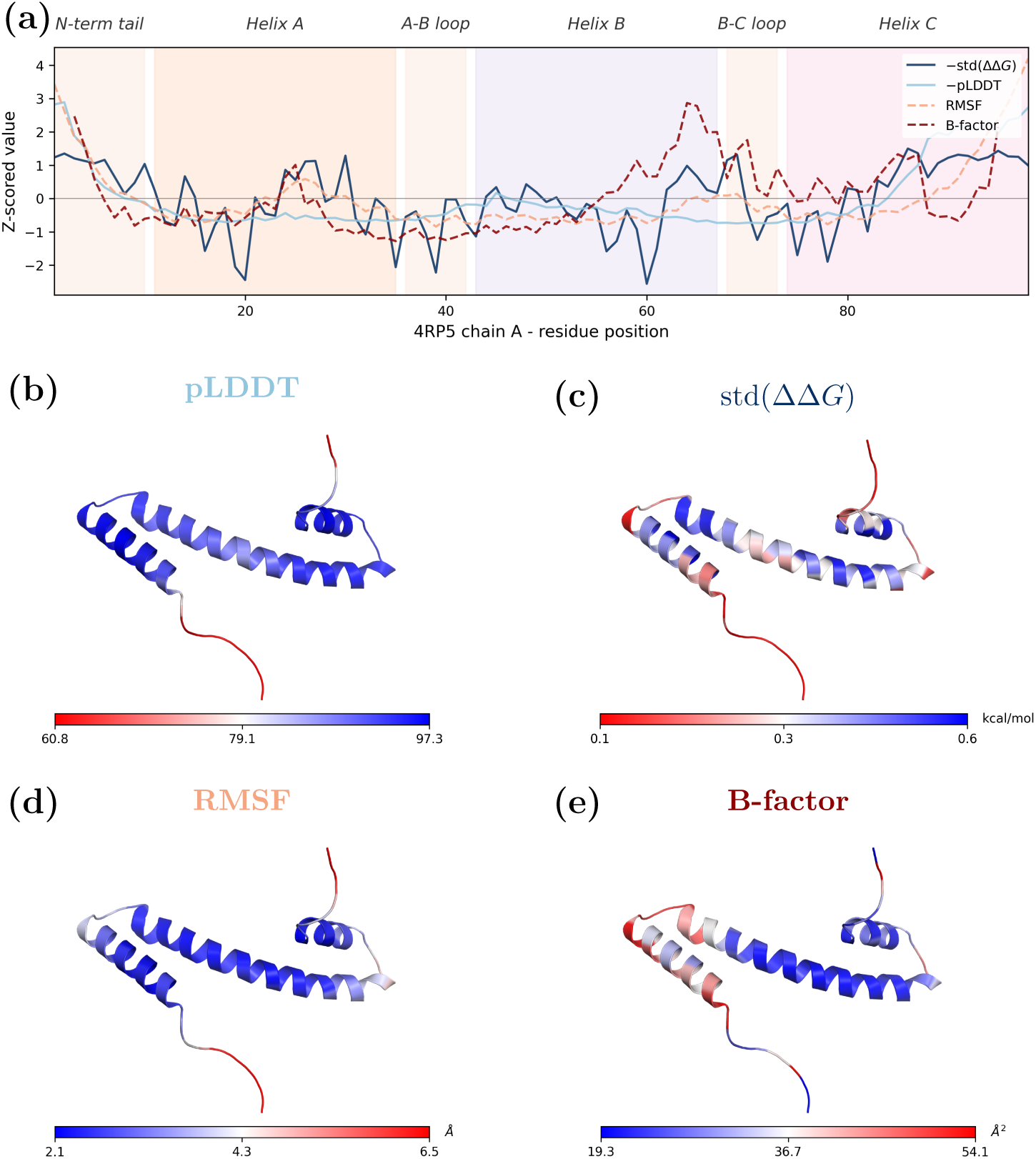
Per-residue robustness vs. dynamics on the L27 domain of *Drosophila* Discs Large 1 (PDB 4RP5, chain A, 98 residues). Layout as in Figure 4. The two-domain helical bundle shows a clear flexible inter-domain linker (red in c and d). pLDDT assigns uniformly high confidence throughout (*ρ*_B-factor_ = +0.01, essentially zero), while robustness captures the flexibility gradient (*ρ*_RMSF_ = −0.78, *ρ*_B-factor_ = −0.33).

**Figure S4:**
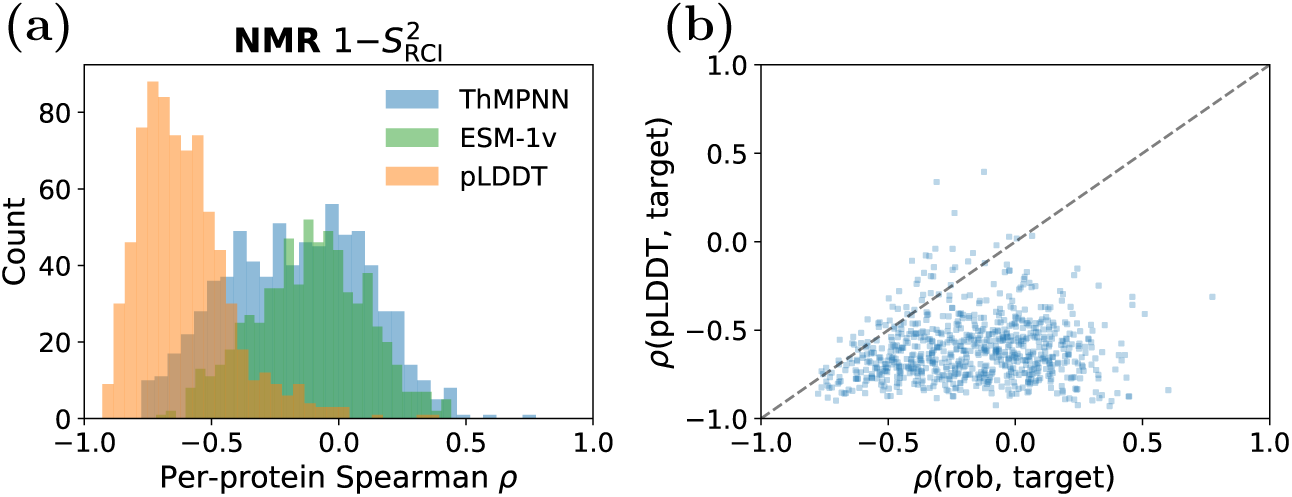
Per-protein Spearman correlation distribution for NMR order parameter (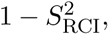 *n* = 759 proteins). Layout as in Figure 1. The robustness distribution is markedly broader than for MD-based targets, with 31% of proteins showing positive (wrong-sign) *ρ*.

**Figure S5:**
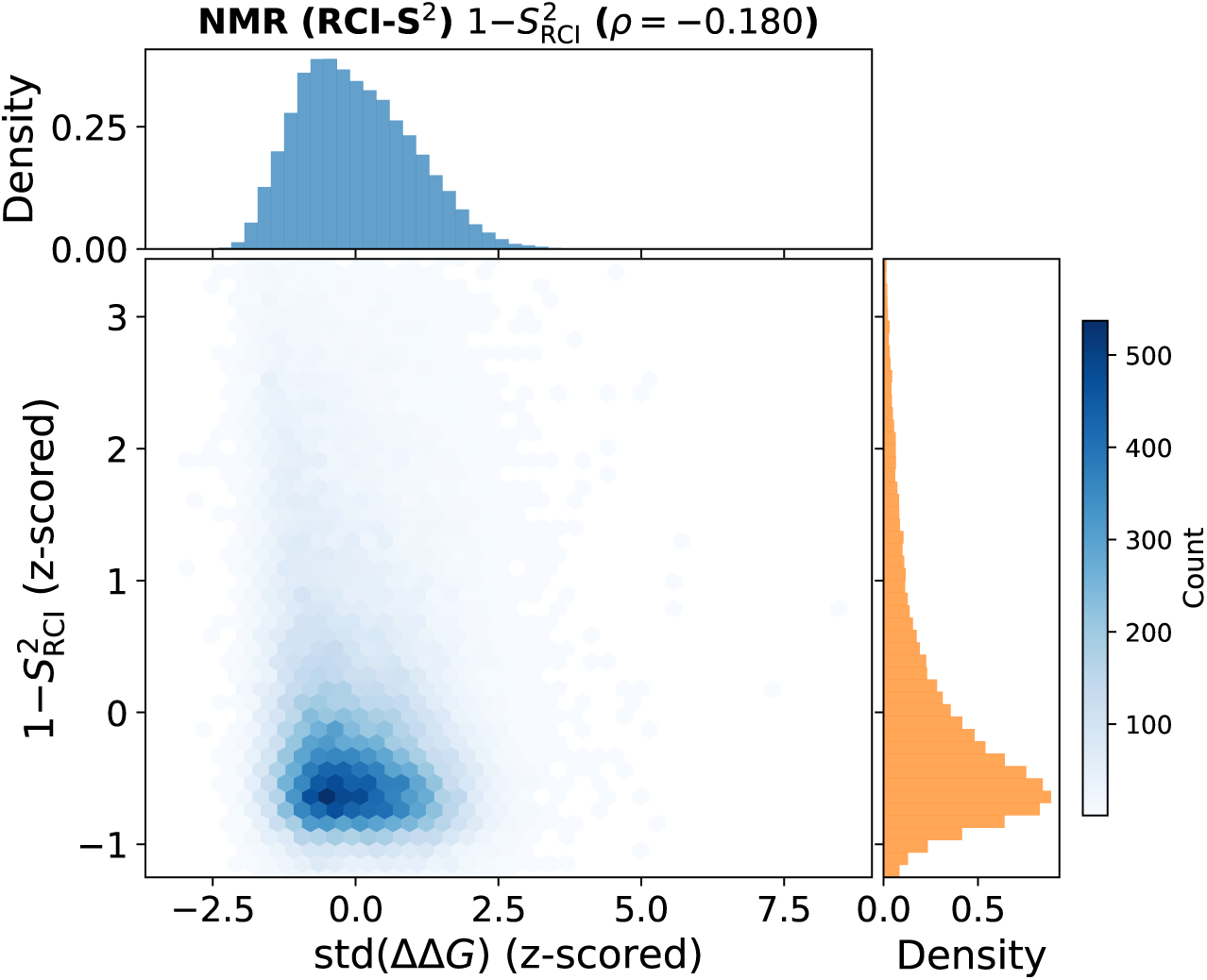
Pooled 2D density of per-residue std(ΔΔ*G*) (z-scored) vs. 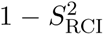 (z-scored) for ThermoMPNN. Layout as in Figure 2. The weaker correlation (*ρ* = −0.176) reflects the distinct nature of NMR-derived order parameters compared to MD RMSF and crystallographic B-factors.

**Figure S6:**
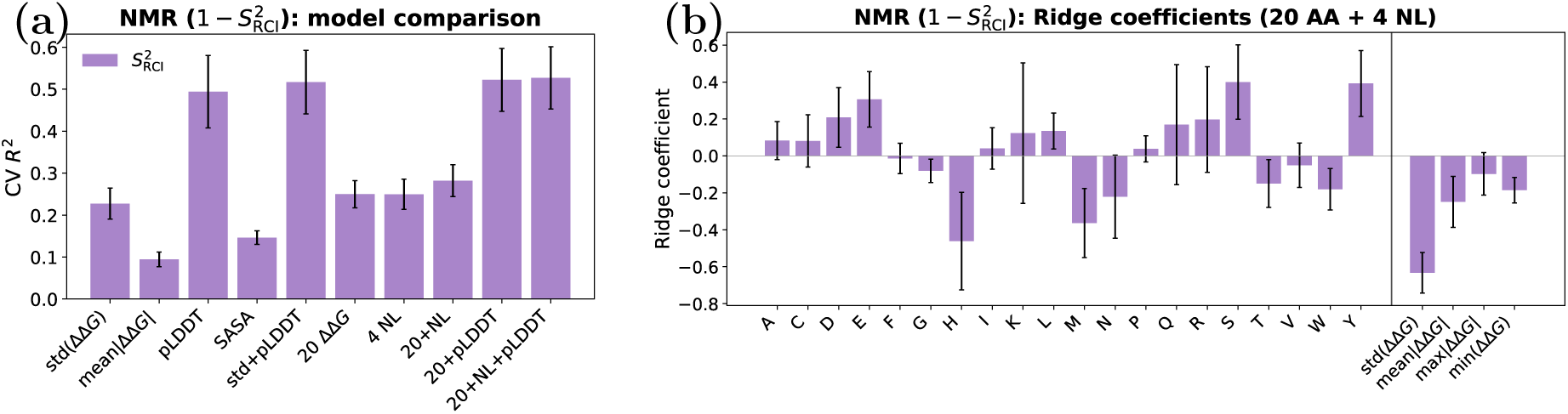
Multi-ΔΔ*G* regression for NMR order parameter 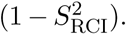 Layout as in Figure 3. **(a)** Cross-validated *R*^2^. **(b)** Ridge coefficients (20 AA + 4 NL features). std(ΔΔ*G*) dominates (−0.63), with per-AA coefficients playing a secondary role; the model achieves CV *R*^2^ = 0.282.

